# Optimized Plaque Assay for Detecting Chronically Infecting Viruses of Haloarchaea

**DOI:** 10.1101/2024.11.12.619373

**Authors:** Sharon Navok, Lee Cohen, Eliora Z. Ron, Uri Gophna

## Abstract

Archaeal viruses predominantly exhibit a chronic lifestyle, where viral particles are released without causing host cell death. Therefore, conventional plaque assays, which are well-suited for identifying lytic viruses, usually fail to detect chronically infecting viruses due to their non-lytic nature. To address this limitation, we developed an optimized plaque assay protocol for detecting chronically infecting viruses of haloarchaea, particularly species within the *Haloferax* genus. By enhancing viral diffusibility and infectivity through adjustments in agar concentration and lowering incubation temperature, this modified protocol improved plaque formation, enabling the detection of viruses that cause mild growth delays. We demonstrate successful plaque formation for two chronically infecting viruses, *Haloferax volcanii* pleomorphic virus 1 (HFPV-1) and lemon-shaped virus 48N (LSV-48N), on representative *Haloferax* strains and highlight the potential for isolating novel viruses from environmental samples. This improved plaque assay is an effective method for the detection and quantification of chronically infecting archaeal viruses, providing a new tool for discovering new viral families in extreme environments.

## Introduction

Bacterial viruses, known as bacteriophages, primarily exhibit two types of lifecycles: lytic and lysogenic. In the lytic cycle, the phage immediately kills and lyses the host cell following an infection. In the lysogenic cycle, the phage integrates its genetic material into the host genome or maintains it as an episome, in both cases remaining dormant until triggered by a cellular stress to enter the lytic cycle.

The plaque assay is a well-established and accurate method for quantifying virus particles produced by an organism, achieved through the observation of plaque formation on an agar overlay of an indicator strain. This technique is also utilized for the isolation of novel viruses from environmental samples by direct plating and facilitates the differentiation of various viruses based on distinct plaque morphologies. In bacteria, clear plaques are typically produced by lytic phages, while turbid plaques are formed primarily by temperate phages, such as coliphage lambda. The turbid plaques result from the presence of a substantial fraction of cells that have become lysogens and are therefore immune to the spread of the lytic phage. Additionally, turbid plaques can also result from phages that produce chronic productive infection, where viruses are continuously released without lysing the host, as in the case of M13 phage and other *Inoviridae*^1^.

Archaeal viruses exhibit considerable morphologic and genetic diversity and most of them possess dsDNA genomes and unique structural properties^2 3 4^. In archaeal viruses all of the three life cycles described above for bacteriophages exist: they can have lytic and lysogenic lifestyle, as well as chronic productive infection^5^. The latter lifestyle has been shown to be much more common in archaea than in bacteria, especially in extreme environments, such as hypersaline ecosystems and acidic hot springs^5^.

To date, traditional methods that rely on plaque formation failed to identify chronically infecting archaeal viruses, shown by other means to be able to infect new hosts^6,7^. In order to obtain plaques, we studied several conditions which could affect plaque formation. Successful plaque formation of viruses requires consideration of several factors, including the host, the virus life cycle, and the growth conditions. In the case of non-lytic lifecycles, plaques are formed through the growth delay of the host, and their morphology, clearness and plaque size are influenced by the ability of virus particles to transfer from cell to cell within the agar overlay (diffusibility and infectivity), as well as the level of growth retardation the virus causes. For better observation of plaques, a significant increase in diffusibility may enhance plaque formation, especially for viruses that cause relatively mild growth delay. Lowering the concentration of agar can increase diffusibility, thereby improving plaque visibility.

In this study, we optimized a plaque assay protocol to facilitate the formation of plaques by chronically infecting haloarchaeal viruses. We enhanced virus diffusibility and infectivity by reducing the agar concentration in the agar overlay and lowering the incubation temperature to slow the growth rate of the haloarchaea.

Our model organisms were species of *Haloferax*, a well-studied archaeal genus that is easy to cultivate. We show that two chronic-infecting viruses isolated from *Haloferax volcanii* strains: *Haloferax volcanii* pleomorphic virus 1 (HFPV-1)^7^ and lemon-shaped virus 48N (LSV-48N)^8^, form plaques on representative haloarchaea strains. Additionally, we demonstrate that this optimized protocol can be used for the isolation of novel viruses from various environmental sources, such as saltwater sinkholes.

## Materials and Methods

### Virus Isolation

Viruses were isolated and purified with PEG (polyethylene glycol) precipitation as detailed in ^7^. In brief, enrichments of archaeal cultures were centrifuged, and the supernatants were precipitated with PEG 8000 overnight at 4°C. The viral particles were subsequently collected by centrifugation at 13,000 g and resuspended in 18% buffered salt water (BSW).

### Archaeal strains and growth conditions

Archaea were grown in Hv-YPC medium ^9^ with shaking at 45°C. For agar plates, 1.5% agar (Difco) was added. For soft agar overlay 0.2% agarose (Lonza) was used. Dilutions were performed using 18% BSW. To start experiments archaea were grown overnight in liquid medium at 45°C with shaking.

### Plaque assay

To prepare the plates, soft agar was prewarmed and cooled to 62°C. Then, 400 µl of overnight cultures (OD∼1/ late exponential growth phase) and 18 µl of 0.5 M CaCl2 were added to 3 ml of 0.2% soft agar mixed well and poured over agar base plates. 3 µl of the virus suspension and serial dilutions were applied to the plates after they solidified. The plates were then incubated at 30°C for 48 hours and left at 25°C for several days until plaques were more easily visible.

## Results and discussion

Our goal was to develop a sensitive plaque assay suitable for the detection and quantification of chronic infection by haloarchaeal viruses. As a proof of concept, we used the optimized protocol described above (Methods) to determine the infectivity of the chronic viruses HFPV-1 (a pleomorphic virus, unable to integrate into the host genome) and LSV-48N (a lemon-shaped virus, which causes chronic productive infection while also integrating into the host genome). As potential hosts, we used six representative strains of *Haloferax*: *H. volcanii* DS2, *H. volcanii* 48N ^10^, *H. volcanii* LSV-cured 48N ^8^, *H. lucentense, H. mediterranei* and *H. gibbonsii* (Table 1).

**Table 1.** Strains and viruses used in this study.

|  | Strain or virus | Reference |
| --- | --- | --- |
| 1 | *H. volcanii* DS2 | 15 |
| 2 | *H. volcanii* 48N | 10 |
| 3 | *H. volcanii* 48N-LSV-cured | 8 |
| 4 | *H. lucentense* Aa 2.2 | 16 |
| 5 | *H. gibbonsii* LR2-5 | 17 |
| 6 | *H. mediterranei* R-4 | 18 |
| 7 | *Haloferax volcanii* pleomorphic virus 1 (HFPV-1) | 7 |
| 8 | Lemon-shaped virus 48N (LSV-48N) | 8 |

### Plaque formation of HFPV-1

Plaques of HFPV-1 were obtained on four of the selected strains: *H. volcanii* DS2, 48N, LSV-cured 48N, and *H. lucentense*, but not on *H. gibbonsii* and *H. mediterranei*. Plaques appeared following 48 hours of incubation at 30°C and plates were left on the bench until more prominent plaques emerged. Importantly, some strains did not exhibit plaques when grown at 37°C, indicating a physiological difference in either the virus or the host that enables better infection at lower temperatures. The most noticeable plaques were observed on

*H. volcanii* DS2, indicating significant growth delay, consistent with its known susceptibility to HFPV-1 infection^7^. In contrast, the virus produced turbid plaques on the other strains, indicating lower growth delay. Curiously, HFPV-1 formed more prominent plaques on 48N-LSV-cured compared to 48N. Infection of 48N by LSV-48N was shown to downregulate several defense systems, and so one would expect that it would be more susceptible to (co-)infection by HFPV-1 due to this “immune-compromised” state. Yet, one should keep in mind that 48N-LSV-cured grows much faster and potentially suffers a more severe (relative) growth delay, since one or more of its defense systems, such as CBASS, may actually cause some cells to enter dormancy or experience growth delay^11^. In some cases, we were able to observe single plaques, allowing accurate determination of infection titer, which varied among the host strains (Table 2).

**Table 2.**
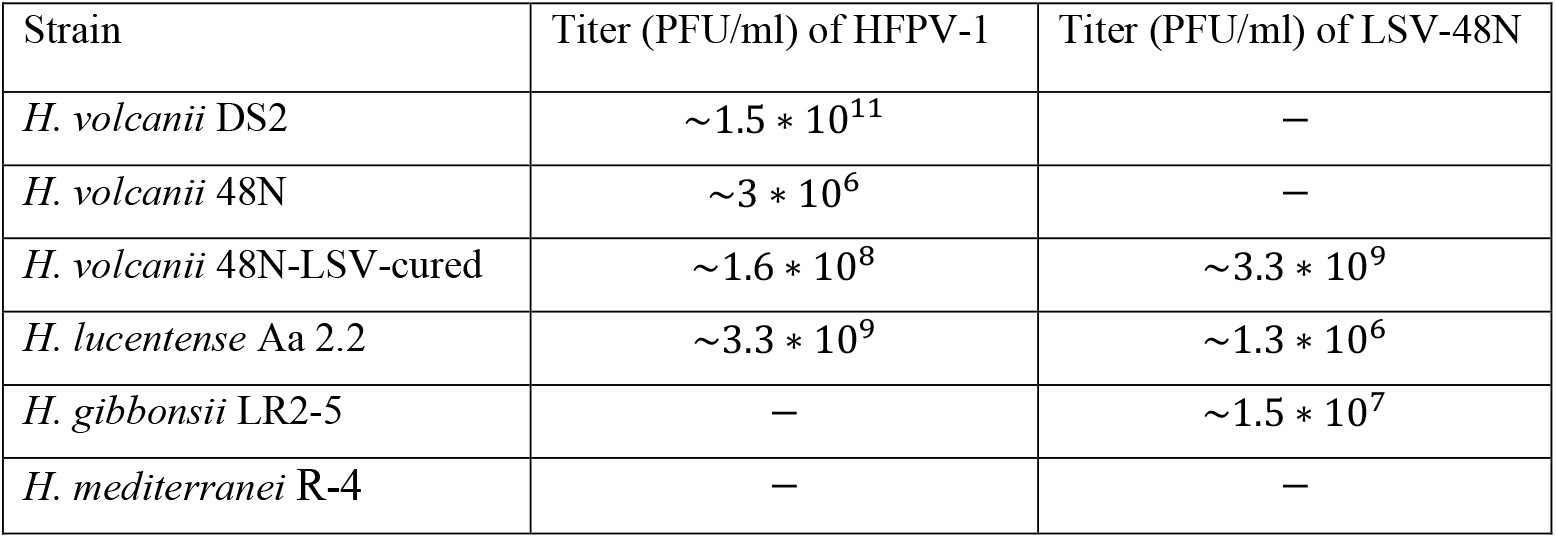
Plaquing efficiency of HFPV-1 and LSV-48N on the representative strains. Virus titer was determined by this formula: 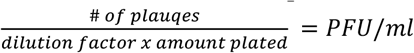

### Plaque formation of LSV-48N

In contrast to HFPV-1, which is known to be a broad-host range virus, plaques of LSV-48N were mainly obtained on 48N-LSV-cured, where it induced clear plaque formation up to the 10^−5^ dilution. For *H. lucentense* and *H. gibbonsii* turbid plaques were only obtained at higher titers (Table 2). As expected, applying LSV-48N to wild-type 48N cells that are already infected did not result in plaque formation, since more viruses of the same species as ones already replicating in the cells are unlikely to cause additional growth delay.

### Isolation of novel viruses

To assess the applicability of this protocol for the isolation of novel viruses from environmental samples, we tested two samples (S1 and S2) collected from Dead Sea sinkholes on the same set of haloarchaeal strains. The samples, sourced from sequenced saltwater sites where haloarchaea were previously identified using metagenomics, produced turbid plaques only on 48N and LSV-cured 48N, indicating the presence of viral activity. Further characterization is needed to identify and characterize the viruses present in these samples.

## Conclusions

Plaque assays represent a rapid, inexpensive and quantitative means to identify new viruses, and measure their infectivity as well as quantify the level of protection of defense systems such as CRISPR-Cas, DISARM^12^, or restriction modification. Plaque assays are straightforward for lytic haloarchaeal viruses ^13 14^, but it has been noted that chronic-infecting archaeal viruses often do not form plaques using current protocols ^7^. Here we present an improved plaque assay protocol that revealed plaque-formation for two viruses, where previously existing protocols were lacking, and demonstrate the potential for isolating new viruses. We hope that with this new protocol new families of chronic-infecting viruses can be isolated from natural samples for *Haloferax* and other haloarchaeal genera.

## Acknowledgements

We thank Dr. Sarit Avrani from the Department of Evolutionary and Environmental Biology at the University of Haifa, Israel, for generously providing the environmental samples used in this study.

---

**Figure 1.**
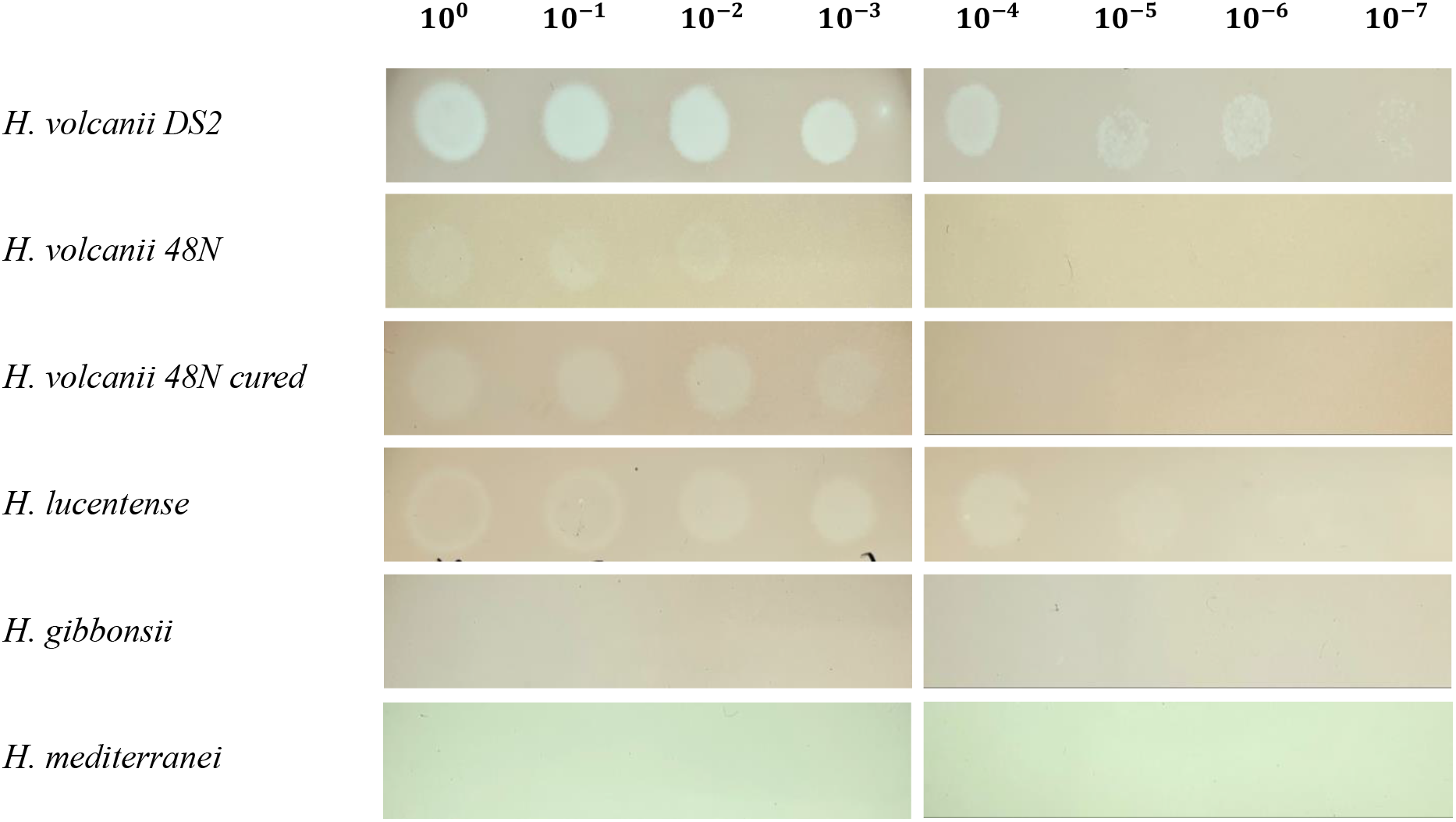
Plaquing efficiency of HFPV-1 on representative haloarchaea strains. Dilutions of virus were plated on soft agar lawns poured on base plates and incubated at 30°C for 48 h, followed by incubation at 25°C to promote more prominent plaque formation.

**Figure 2.**
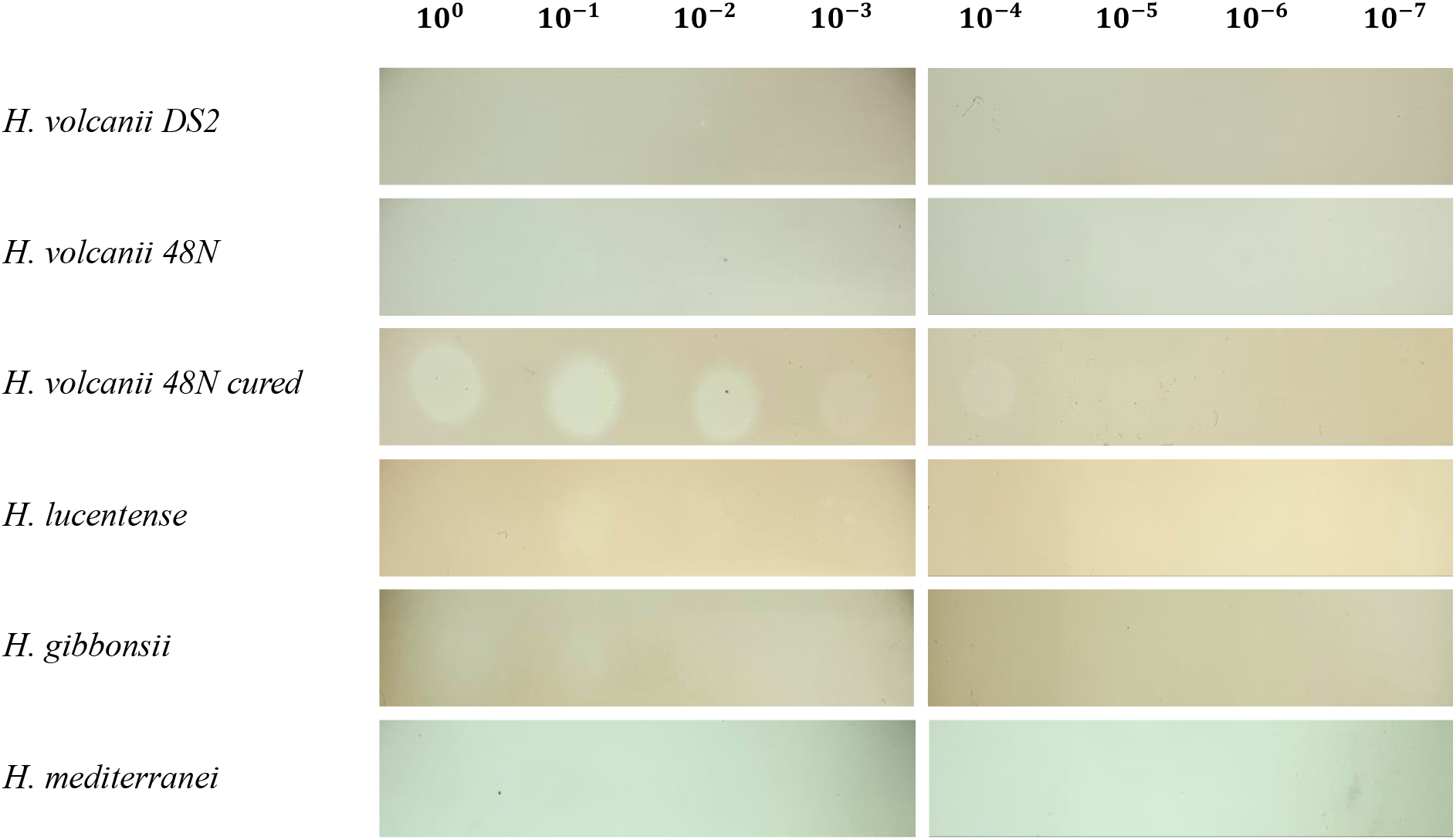
Plaquing efficiency of LSV-48 on representative haloarchaea strains. Dilutions of virus were plated on soft agar lawns poured on base plates and incubated at 30°C for 48 h, followed by incubation at 25°C to promote more prominent plaque formation.

**Figure 3.**
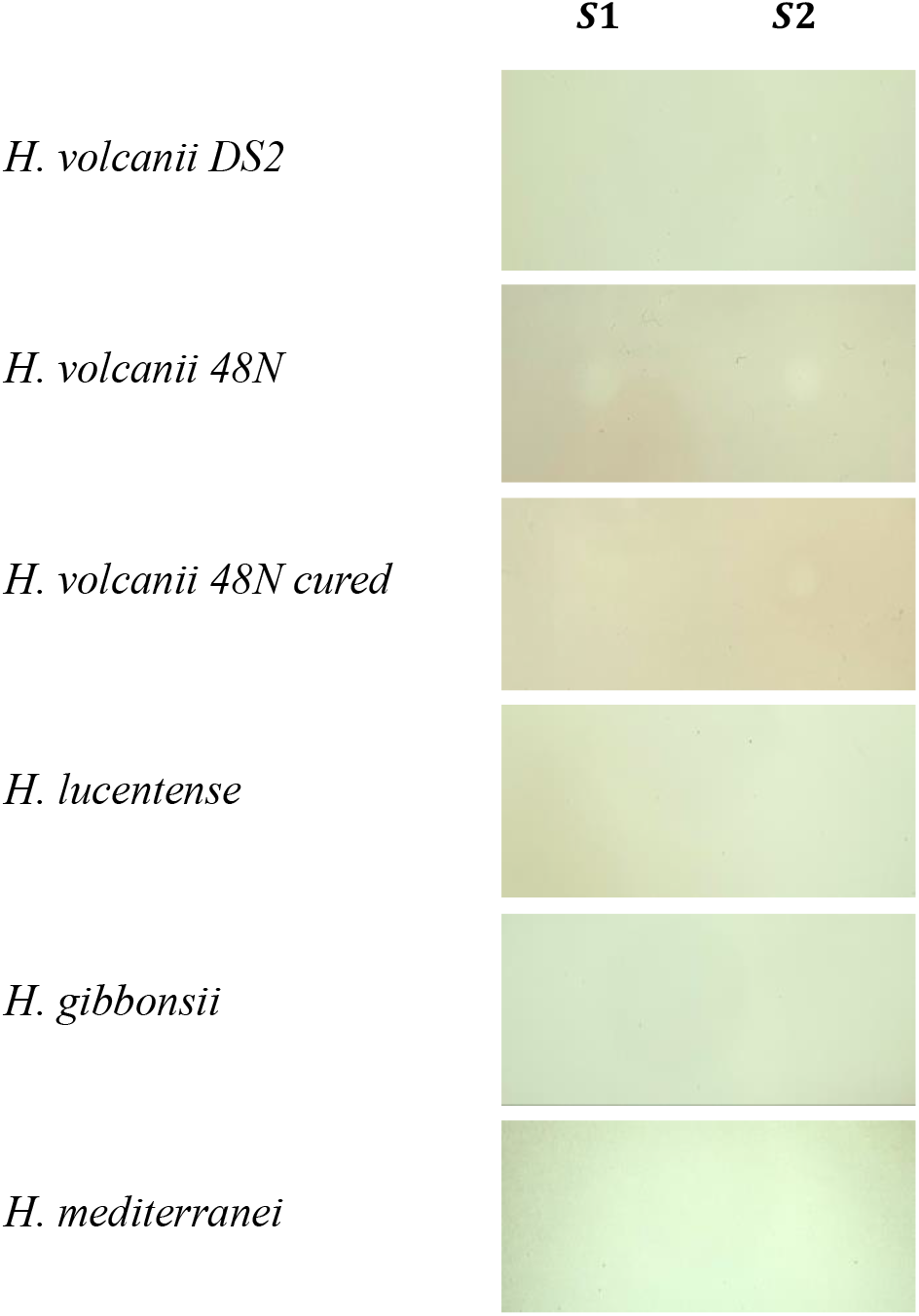
Plaque formation of environmental samples on representative haloarchaea strains. Two environmental samples (S1 and S2) were plated on soft agar lawns poured on base plates and incubated at 30°C for 48 h, followed by incubation at 25°C to promote more prominent plaque formation.

## Notes

### Competing Interest Statement

The authors have declared no competing interest.

